# Differentiation hierarchy in adult B cell acute lymphoblastic leukemia at clonal resolution

**DOI:** 10.1101/2025.09.10.674954

**Authors:** Alec Geßner, Adrien Jolly, Tjeerd P. Sijmonsma, Yves Matthess, Manuel Kaulich, Stefan Günther, Fabian Lang, Florian Buettner, Michael A. Rieger

## Abstract

While a differentiation hierarchy with leukemia-initiating stem cells (LICs) at the apex is well documented for acute myeloid leukemia, the existence of LICs and their trajectories in B cell acute lymphoblastic leukemia (B-ALL) are debated. B-ALL is a malignant disease displaying considerable phenotypical and functional heterogeneity, yet the underlying cellular organization remains largely elusive. This study aims to investigate the hierarchical landscape of B-ALL by combining barcoding and multiome single cell data to unveil the differentiation architecture and temporal dynamics of leukemic differentiation at clonal and single cell resolution *in vivo*.

Single-cell transcriptome and AbSeq analysis of lentiviral barcoded and transplanted patient-derived B-ALL cells revealed interconnected subpopulations, which could be prospectively isolated via surface markers and exhibited distinct leukemogenicity. Barcode tracking demonstrated the ability of cells to differentiate from a source cluster into differentiated progeny. Using machine learning we identified expression patterns predicting their differentiation potential of each barcoded cell within the immature cell compartment. To determine the dynamical properties of clones, we have designed a mathematical model that simulates the development of these subpopulations from a clonally diverse stem cell compartment. The model confirmed the most likely cluster of origin, supporting our functional data, and revealed the variability in dynamical properties between clones, while largely excluding plasticity. Hence, our work demonstrates a unidirectional differentiation in B-ALL with an immature population exhibiting a high leukemogenic potential.

**Key points:**

- A differentiation hierarchy with distinct leukemogenesis potential in B-ALL is demonstrated via clonal tracing and mathematical modeling.
- Clonal differentiation behavior, while heterogeneous, is cell intrinsically controlled and predictable, yet independent from expansion.

## Introduction

Tumor cell heterogeneity at inter- and intra-patient level is caused by clonal evolution induced by private somatic genetic alterations in individual tumor cells and an epigenetic tumor cell differentiation hierarchy^1–3^. The models of clonal evolution and cancer stem cell (CSC) hierarchies are not mutually exclusive^4,5^.

A hierarchical cancer cell development involving differentiation and loss of carcinogenesis has been proposed in breast^6^, brain^7^, and colorectal cancer^8^ among other entities. Particularly, in acute myeloid leukemia (AML), the existence of leukemia-initiating stem cells (LICs) has been first experimentally proven by the seminal experiments of Bonnet and Dick in 1997^9^. LICs are functionally defined by their unique ability to engraft and initiate leukemia in immune-compromised mice, and they comprise therapy-resistant cells causing relapse in patients. While still challenging to find common surface markers for LICs in AML^10–12^, an enrichment of LICs became quantifiable based on gene expression signatures^13^. Hence, these markers can now be used in a clinical context^14^ and LICs represent promising therapeutic targets^3,15^.

In contrast, while genomic, phenotypic, and functional cell heterogeneity is well documented in B cell precursor acute lymphoblastic leukemia (B-ALL), the existence of a differentiation hierarchy with LICs at the apex remains controversial^16–19^. B-ALL is a disease that also affects children, though survival rates are lower in adults. While remission rates after first-line therapy exceed 90 %, the prognosis after relapse is dismal with survival rates under 10 %^20,21^. It has been suggested that low proliferating cell subsets within the leukemic population are associated with therapy resistance^22^, but no *bona fide* stem cell at the top of a leukemic hierarchy has been identified. B-ALL cell surface phenotypes vary widely between and within patients^4^.

While scRNA-Seq technologies resolve cell type identities and cell states, recent advances in lineage tracing now allow the reconstruction of the differentiation trajectory of single cells. Lineage tracing approaches, using genomic and mitochondrial DNA mutations as endogenous barcodes, have suggested the existence of multiple paths of cell differentiation in AML^23^, thereby uncovering a leukemia hierarchy parallel to normal hematopoiesis and the existence of a differentiation bias at clonal level. A few studies used lentiviral genomic barcoding to analyze the clonal composition of infant B-ALL primografts, which had experienced already a strong selection before barcoding the cells^24,25^. By combining the information from cell barcodes expressed at the RNA level that are transferred to progeny throughout generations, scRNA-Seq can infer lineage development in embryogenesis and in adult stem cell systems^26–28^. This type of approach offers a unique chance to learn about differentiation trajectories in B-ALL. In this work, we use cellular barcoding, serial-xenotransplantation, and single-cell sequencing to study the development of adult B-ALL *in vivo* at clonal and single-cell level.

## Methods

### Xenotransplantations

NOD.Cg-Prkdcscid Il2rgtm1Wjl/SzJ (NSG) mice were housed at the animal facility of the Georg-Speyer-Haus (Frankfurt, Germany) under pathogen-free conditions. Transplantations were performed with male and female animals (at 8-12 weeks of age) and littermates were randomly assigned to experimental groups. All experiments were approved by the local authorities (FK-2004, Regierungspräsidium Darmstadt, Germany) and followed the rules of the German animal welfare legislation.

Irradiated (2.5 Gy) NSG mice were intravenously inoculated with either 1×10^6^ (bulk) or 1×10^5^ (sorted subpopulations) barcoded cells via the tail vein. PB analysis was performed every 2-4 weeks post-transplantation by flow cytometry with anti-human CD45-BV711 (clone HI30, BD) and anti-mouse CD45.1-PerCP-Cy5.5 (clone A20, Biolegend). Mice were sacrificed once they reached the stop criteria due to their leukemia burden. BM isolated by flushing femurs and tibiae were stained with antibodies for flow cytometric analysis. For secondary transplantation, 5×10^5^ BM tumor cells were intravenously injected into 2.5 Gy-irradiated NSG.

### Cell culture

Patient CR and BV-derived B-ALL cells (both male) from an early passage were freshly thawed and cultured at a cell density of 1×10^6^ cells/ml in IMDM (Thermo Fisher Scientific)-based medium according to the established protocol^29^. Lenti-X HEK293T cells (#632180, Takara Bio) were cultured in DMEM containing 10% FBS, 1% L-Glutamine and 1% Penicillin-Streptomycin (all from Thermo Fisher Scientific). All cells were cultured at 37 °C with 5% CO_2_ and 95% relative humidity.

### Lentiviral Barcoding

The barcoding plasmid (see supplemental methods) was encapsulated within vesicular stomatitis virus G (VSVG)-pseudotyped lentiviral particles using a split-genome strategy through calcium phosphate transfection of Lenti-X HEK293T cells^30^. Virus particles were enriched via ultracentrifugation at 50,000×g for 1 h at 4 °C and titers were quantified by limited dilution transduction of patient-derived B-ALL cells via flow cytometry (mCherry expression). The barcodes were stably integrated into the genome of patient CR- and BV-derived B-ALL cells by lentiviral transduction with a multiplicity of infection of 0.1.

### Flow cytometry

Assessment of cell viability was conducted with the Fixable Viability Dye eF780 (Thermo Fisher Scientific). Surface marker expression was analyzed at an LSR Fortessa (BD) using antihuman CD44-FITC (clone L178, BD) and anti-human HLA-ABC-V450 (clone G46-2.6, BD). Cells were sorted utilizing a FACS Aria III (BD).

### Single-cell Sequencing

Single cell sequencing was performed with the Rhapsody system (BD) as detailed out in the supplemental methods.

### Single-Cell data analysis

The reads were aligned and counted using BD SevenBridges and then analyzed in R with Bioconductor and Seurat. The machine class prediction was performed with XGboost and the predictions tested with permutation tests (see detailed description in supplemental methods).

### Mathematical modeling of leukemia development

A detailed description of our model and approach to parameter estimation is presented in supplemental data

## Results

### B-cell subpopulations with differentiation states emerge *in vivo*

To combine the analysis of subclonal architecture and differentiation trajectories of individual clones, we created lentiviral barcoding technologies to track LICs and their fate at clonal and single-cell resolution *in vivo*. Adapted from the high-complexity ClonTracer library^31^, providing 4×10^7^ unique base pair sequences (= barcodes), we generated an expression vector that encodes for a mRNA with the coding sequence of the red fluorescent protein mCherry followed by the 30-mer semi-random barcode in the 3’ UTR (**supplemental Figure 1A**). Importantly, the synthesized barcode library was inserted in an equimolar ratio into the vector using the cloning-free 3Cs technology^32^. The lentiviral delivery method facilitates efficient single barcode integration into the host genome, ensuring stable inheritance and barcode expression throughout cell divisions. Finally, the ClonTracer compatibility with next-generation and single-cell sequencing allows for accurate analysis of barcode abundance, providing a quantitative readout of clonal dynamics. Transduced cells showed robust and persistent mCherry expression for more than 6 months, not observing effects of promoter silencing (**supplemental Figures 1B and 1C**). Sequencing data from a transduced library 3 months after transduction revealed an even distribution of most barcodes throughout the library, each appearing at frequencies ranging from 0.025 % to 0.075 % for 3958 cells (**supplemental Figure 1D**).

**Figure 1.**
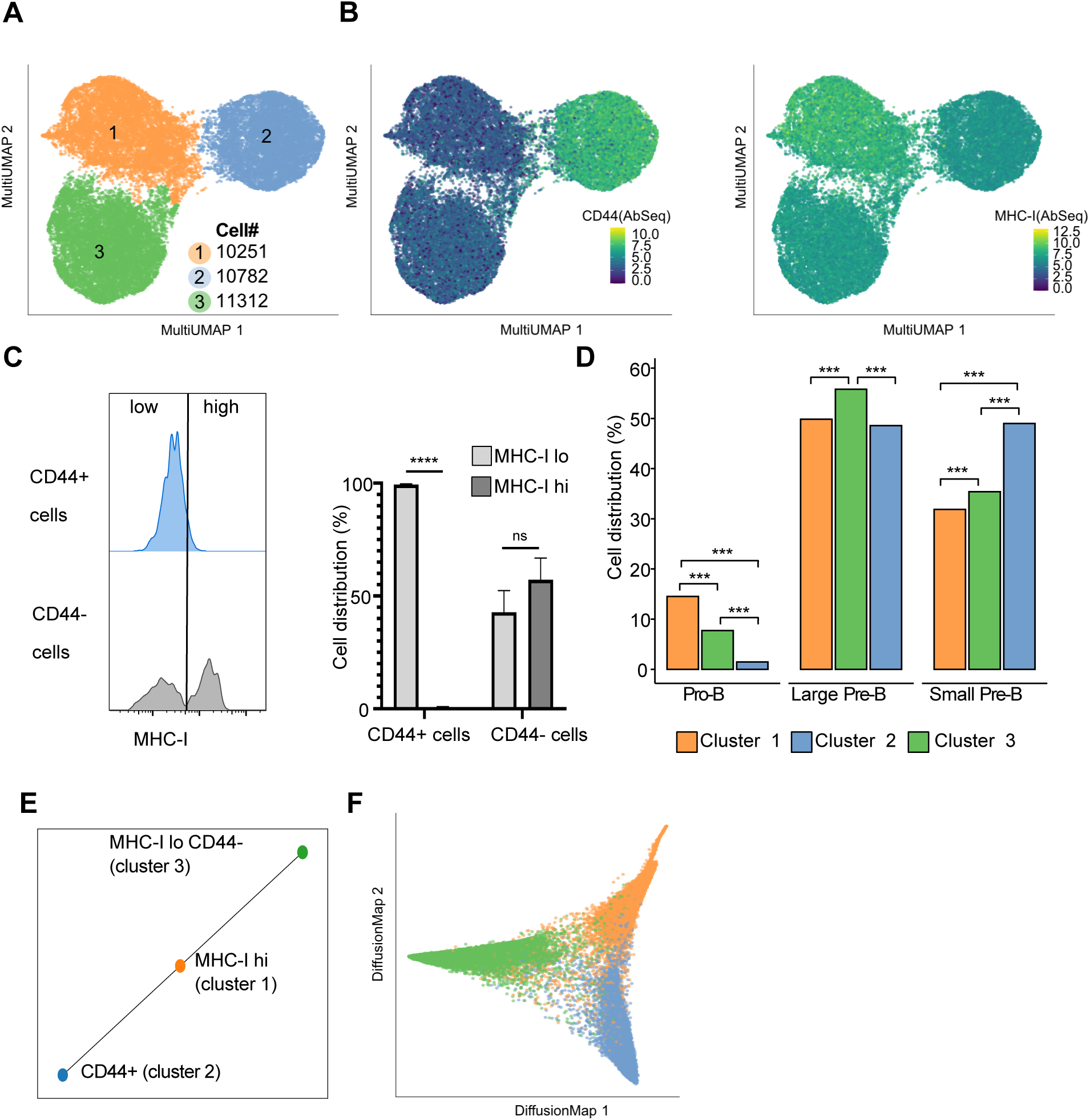
Single-cell expression reveals distinct cell stages. (A) MultiUMAP visualization of mRNA and surface protein expression-based clustering analysis of 35993 leukemic cells isolated from murine BM (n = 2 mice). (B) AbSeq expression of CD44 and MHC-class I. (C) MHC-class I surface expression of CD44+/-patient CR-derived B-ALL cells after transplantation by flow cytometry. Representative histogram and bars represent the mean with SEM (n = 4 mice). Two-way ANOVA with Šídák’s multiple comparisons test. (D) Proportion of cells assigned to the Pro-B, Large Pre-B and Small Pre-B stages of B-cell development per cluster. Pairwise comparison Chi-squared test with Bonferroni correction. (E) Partition-based graph abstraction (PAGA) reveals cluster 1 as the only connection hub between clusters 2 and 3; threshold = 0.1. (F) Diffusion map with colors corresponding to clusters presented in A.

Next, we used the lentiviral particles to transduce patient-derived adult B-ALL cells from the patients CR and BV^29^. *In vivo* leukemogenicity of both samples was confirmed by intravenously injecting 1 × 10^6^ unsorted bulk cells into NSG mice. For patient CR-derived B-ALL, the recipient mice were sacrificed at day 49 due to accelerated tumor burden. Leukemic cells engrafted in BM, PB and spleen (supplemental **Figure 2A**). FACS-sorted BM-derived mCherry^+^ B-ALLs cells were stained with 49 oligo-nucleotide-labeled antibodies (AbSeq) against surface markers (supplemental **Table 1**) and single-cell sequencing was performed using the BD Rhapsody platform. Clustering of cells fromfrom the BM of recipient mice revealed three clusters based on combined mRNA and AbSeq expression, indicating the existence of molecularly distinct populations within the leukemia (**Figure 1A**). Differential surface marker expression analysis of the three clusters based on AbSeq demonstrated a higher expression of MHC-class I on cells of cluster 1 (MHC-I hi), an upregulation of CD44 and a down-regulation of MHC-class I on cells of cluster 2 (referred as CD44+), while cluster 3 (referred as MHC-I lo CD44-) showed low expression in MHC-class I and CD44 (**Figures 1B and supplemental Figure 2B**). The differential surface expression of CD44 and MHC-class I of ALL cells at the stage of full-blown leukemia in NSG mice was confirmed by flow cytometry (**Figure 1C, supplemental Figure 2C-E**). Of note, little differences in cell cycle distribution were observed **(supplemental Figure 2F)**.

As B-ALL reflects aberrant B-cell differentiation, we mapped our dataset to normal B-cell development to evaluate at which B-cell progenitor stage the clusters were mostly associated^33^. We observed that while most cells were assigned to the large Pre-B state, cells from cluster 1 were significantly more associated with earlier developmental stages than the cells of the two other clusters (**Figure 1D**). Cluster 1 showed an over-representation of cells assigned to pro-B cells and an under representation in the small pre-B stage. We investigated the connectivity of the three cell clusters by PAGA^34^ analysis and found that cluster 1 cells comprise the connecting hub of cells from cluster 2 and cluster 3 (**Figure 1E**). To arrange cells according to their position on the differentiation trajectory, we performed a diffusion map ^35,36^, which indicated that cluster 1 lies at the tip of the differentiation system (**Figure 1F**). Taken together, these data suggest that cells in cluster 1 comprise a less differentiated state of B-ALL cells and that cells in clusters 2 and 3 are directly related to cluster 1 cells and represent more differentiated cell populations.

Likewise, we could identify 2 distinct sub-populations in the B-ALL dataset derived from patient BV, reflecting distinct stages of B-cell development pointing to differentiation with cluster 1 being cluster of origin (supplemental **Figures 3A-B**) and notably characterized by differential surface expression of CD38 (supplemental **Figure 3D**).

### Clonal tracing reveals distinct differentiation behaviors

Next, we focused our analysis on tracking individual clones comprising cells with identical barcodes. From 12867 barcoded cells, we found 118 clones, ranging in size from 2 to 2299 cells (**Figure 2B**). As exemplified by the eight largest clones in our dataset shown in **Figure 2A**, we found considerable heterogeneity in the distribution of clonal cells in the three clusters. While cells of a clone tended to accumulate in a specific cluster, we noted that cells of the same clone were present in multiple clusters indicating that they can change their molecular program. Importantly, while the 10 largest clones comprised 84% of all the barcoded cells (**Figure 2B**). Since we did not observe clonal dominance in culturing the cells *in vitro* for three months (supplemental **Figure 1D**), the microenvironment may strongly influence the expansion of particular clones. Next, we investigated the distribution of cells from each of the 118 clones across the three clusters. Cluster 1 showed the highest diversity with 71 % of the clones having cells in cluster 1, whereas clusters 2 and 3 contain cells from 53 % and 39 % of the clones respectively (**Figure 2C**). Consistent with our findings in Figure 1, the higher clonal diversity suggests that cluster 1 may comprise the source cluster of B-ALL differentiation in this patient.

**Figure 2.**
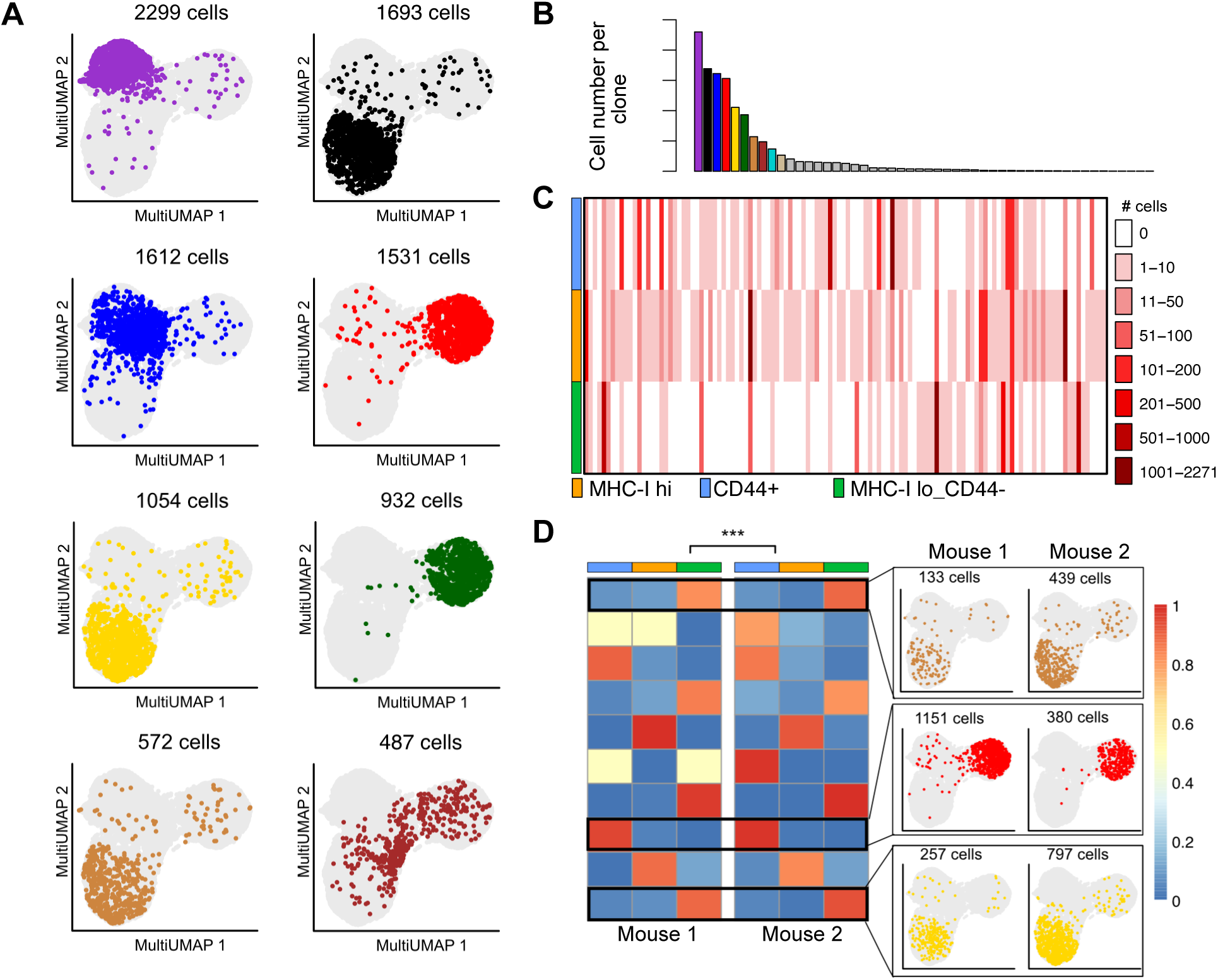
High clonal diversity in the most immature B-ALL cluster. (A) UMAP projections display the localization of the eight largest clones. (B) Clone size distribution. (C) Clonal diversity in the clusters. Each line represents a clone, colored by clone identity. (D) Heatmap depicting the distribution of shared clones across the three clusters in both mice. P-value of similarity determined by permutation test (see supplemental methods). **, p≤0.01; ***, p≤0.001.

We considered the distribution of cells in clusters 1, 2, and 3 of each clone to analyze distinct differentiation patterns and behavior. This analysis revealed that identical clones overlapping between replicate mice (**Figures 2D and supplemental 4A**) showed highly similar differentiation patterns across recipients, suggesting a cell-intrinsic differentiation behavior. Importantly, this similarity contrasted with the size of these clones (supplemental **Figure 4B**) which showed no correlation between replicates. In conclusion, while differentiation potential is cell intrinsic in these clones, clone growth may depend highly on the local environment.

Next, we performed a PCA on the clone distribution across clusters, thereby revealing three distinct classes of clones highly separated by the two first components(**Figure 3A**). These three classes were characteristic for cells to either differentiate in cluster 2 (cluster-2-biased class = class B) or cluster 3 (cluster-3-biased class = class C), or to remain immature in a self-renewal-like stage (self-renewing class = class A).

**Figure 3.**
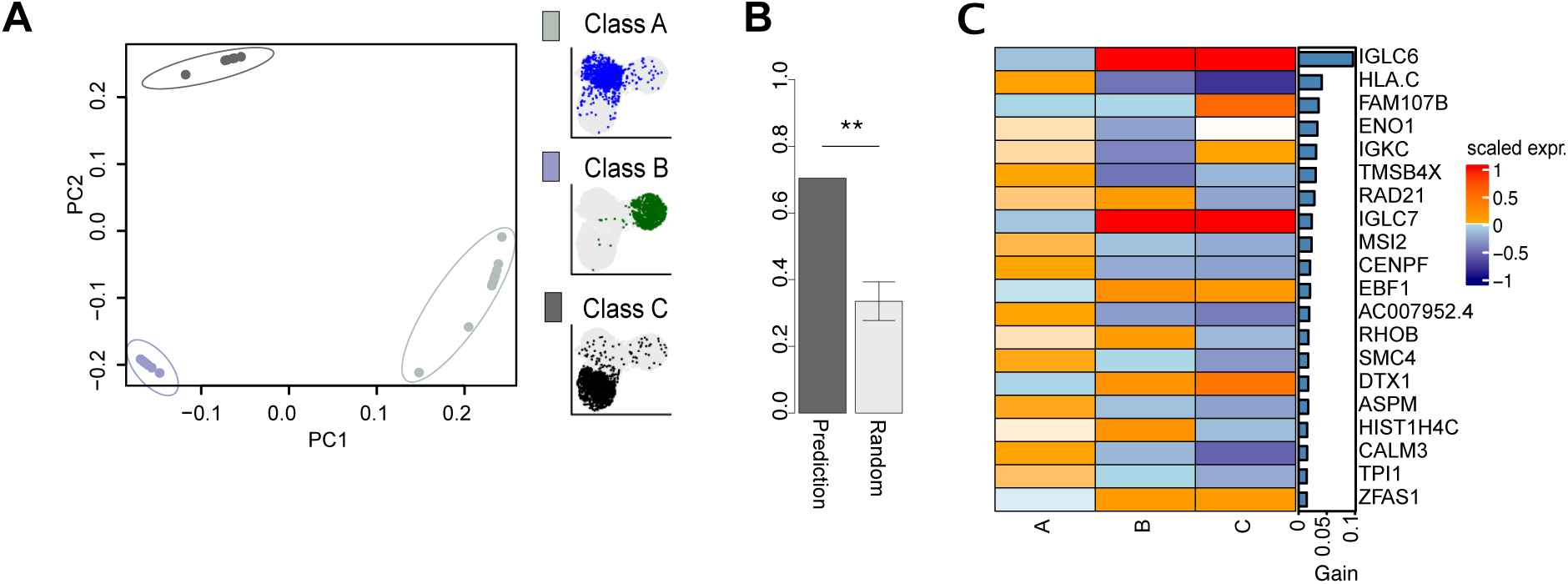
Predicting differentiation classes. (A) PCA of all clones with more than 20 cells identifies three classes of differentiation potential. (B) Balanced accuracy of the XGboost model versus mean balanced accuracy from 1000 label permutations in the training set (see supplemental methods); error bar denotes SD. (C) Top 20 class predictors ranked based on their prediction gain with their relative average expression (z-score) in each class in cluster 1.

Since cells in cluster 1 represent progenitors of cells differentiating into clusters 2 and 3, we hypothesized that the transcriptome of cells in cluster 1 was sufficient to predict the differentiation behavior and future fate of their progeny. We therefore quantified to what extent the transcriptome of a cell in the source cluster was already indicative of a differentiation bias. To this end, we used XGBoost^37^ with 90 % of the clones as a training set supplemental Methods, to identify genes that drive the decision of cells to remain self-renewing (class A) or belong to the differentiation-biased classes B or C. With this trained model, we obtained a balanced accuracy of 0.7 in predicting the differentiation outcome of the cells in our test set (**Figures 3B and supplemental Figure 4C**). A permutation test revealed that this accuracy was significantly higher than random predictions, showing the transcriptome of a cell in the cluster of origin was indeed predictive of its differentiation potential. We represented the relative average transcriptome of the 20 most important predictive genes in each class (**Figure 3C**), and noted that e.g. Musashi2 (MSI2), a key regulator of self-renewal program in cancer stem cells^38^, was enriched in the self-renewing class A while EBF1^39^, a key driver of B cell differentiation, had a lower expression in this class. These elements support our claim that class A is a low differentiating class. In the same manner, we found two classes of clones in patient BV-derived B-ALL cells based on their cluster distribution (**supplemental Figures 3C, 3E and 3F**) and could identify markers of differentiation potential within the prospective cluster of origin (**supplemental Figures 3G-H**).

### Mature B-ALL subpopulation exhibits lower leukemogenic potential

Next, we sought to functionally confirm that the three clusters 1, 2 and 3 represent B-ALL cells at different differentiation levels. Hence, we tested their leukemogenic ability by serial xenotransplantations *in vivo*. Our data so far suggested that CD44^+^ CR patient-derived cells (cluster 2) represent a more mature B-ALL subpopulation with lower leukemia-initiating potential compared to CD44^-^ cells. Therefore, we transplanted prospectively FACS-sorted CD44^+^ and CD44^-^ cells in sublethally-irradiated NSG mice (**Figure 4A**). As shown in **supplemental Figure 5A**, 4.2 % of cells showed high CD44 surface expression before transplantation. Each recipient received 50,000 FACS-sorted CD44^+^, CD44^-^ or parental cells, and leukemogenesis was monitored in the PB by flow cytometry. While the median survival of CD44^-^ cell-receiving mice was 75 days after transplantation, the survival of mice receiving CD44^+^ B-ALL cells was substantially longer (154 days median survival), and two out of five mice did not develop leukemia at all (**Figure 4B**). These striking differences functionally demonstrate reduced leukemogenic potential of CD44^+^ cells with delayed measurable disease onset and significantly prolonged survival (p<0.01) of the recipients than the CD44^-^ ALL population and establishing that some cells completely lack leukemogenesis potential.

**Figure 4.**
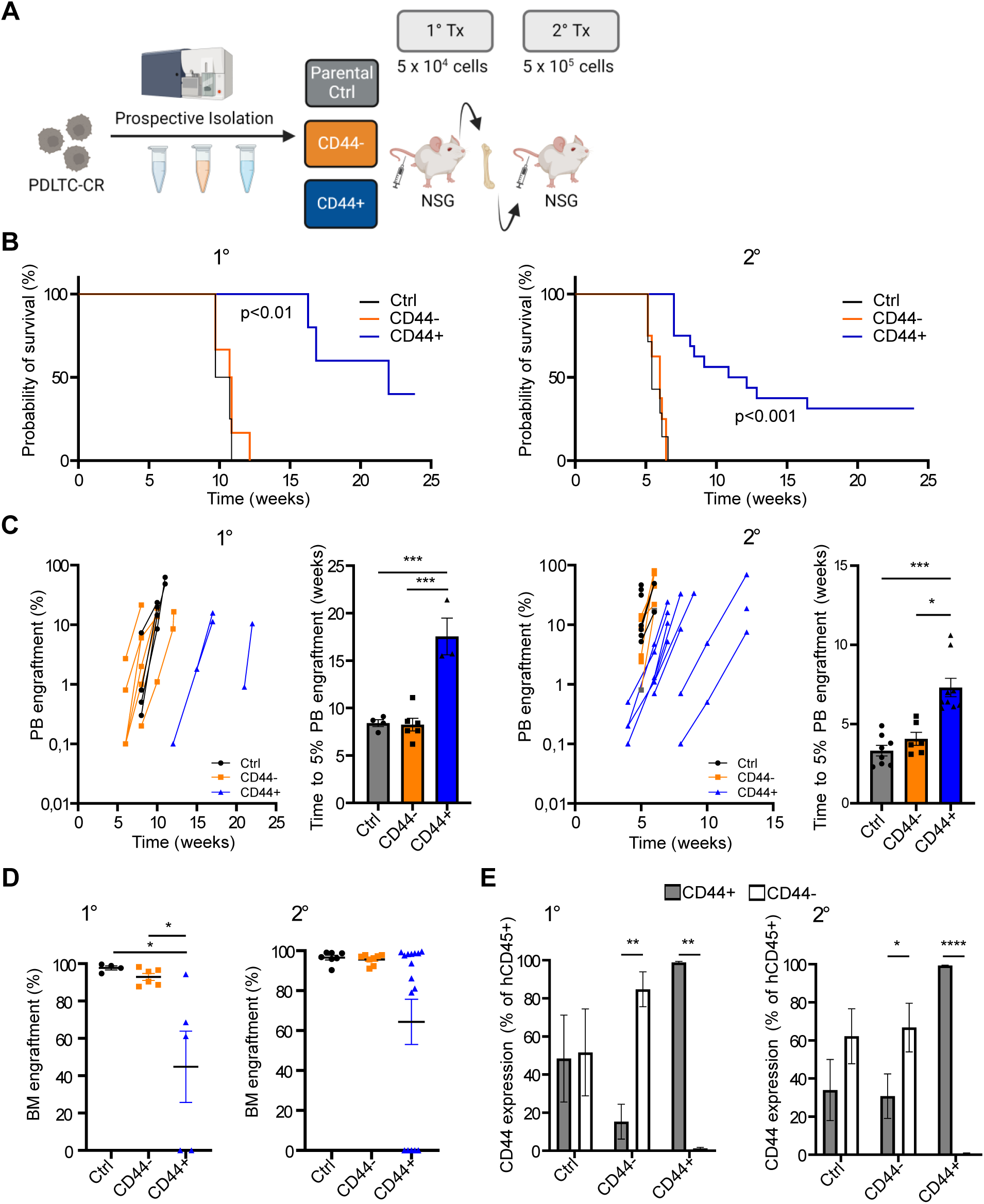
Transplantation of prospective cell stages confirms distinct leukemogenesis. (A) Experimental overview. (B) Kaplan-Meier survival analysis reveals higher survival in mice transplanted with CD44^+^ cells. 1°: n=4-6 mice, 2°: n=7-8 mice. Mantel-Cox test adjusted with Holm-Šídák analysis. (C) Engraftment progression in PB and time to 5 % PB engraftment. Bars represent the mean with SEM. Kruskal-Wallis test with Dunn’s multiple comparisons test. (D) Engraftment levels in BM at the endpoint. Individual values with the mean ± SEM. 1°: One-way ANOVA with Tukey’s multiple comparisons test. 2°: Kruskal-Wallis test with Dunn’s multiple comparisons test. (E) Percentage of CD44^+^ cells in BM. Bars represent the mean with SEM. Two-way ANOVA with Šídák’s multiple comparisons test. Only pairwise comparisons with a significant difference are shown. *, p≤0.05; **, p≤0.01; ***, p≤0.001; ****, p≤0.0001.

We then isolated B-ALL cells from the BM of primary recipients and transplanted 500,000 cells into secondary recipients. Of note, we only transplanted B-ALL cells from mice with leukemia development. While secondary recipient mice receiving B-ALL cells from CD44^-^ and parental cell-transplanted mice succumbed already after 38-42 days, mice receiving cells from CD44^+^ B-ALL-transplanted mice showed a much prolonged survival (p<0.01) or showed no leukemia development at all (**Figure 4B**). These functional tests clearly establish the notion that some cells completely lack leukemogenesis potential.

Specifically, the first detection of B-ALL cells in the PB was at 42 days and 84 days after transplantation in mice receiving CD44^-^ and CD44^+^ cells, respectively (**Figure 4C**). Overall, the increase in leukemic burden in the PB over time was comparable between CD44^+^ and CD44^-^ mice suggesting that the delayed onset in CD44^+^ ALL cells resulted from a markedly reduced frequency of leukemia-initiating cells, however, with a similar proliferation as the parental or CD44^-^ cells. The CD44^+^ transplanted group showed lower engraftment levels of blasts in BM (**Figure 2D**). Importantly, CD44^+^ surface expression remained constantly high throughout disease development, suggesting that CD44^+^ cells maintained their phenotype and did not transition to a CD44^-^ state (**Figure 4E**). Since the growth rate of both subpopulations is similar (**supplemental Figure 5B**), the reason may well be the lower leukemia-initiating potential of CD44^+^ cells. Bulk RNA-Seq of sorted CD44^+^ and CD44^-^ ALL cells from murine BM at full-blown leukemia stage revealed significant differences in gene expression profiles. KEGG pathway analysis demonstrated increased B-cell receptor signaling in CD44^+^ cells (**supplemental Figure 6**). In conclusion, these data functionally confirm the decreased capability of CD44^+^ mature B-cell ALL subpopulation in promoting disease progression and re-emergence in NSG mice.

### Mathematical modeling identifies unidirectional differentiation as the most likely development dynamics

We next aimed at quantifying the dynamics of disease development and learning the clonal differentiation behavior and proliferation characteristics. Since clonal differentiation dynamic cannot be observed directly, mathematical modeling approaches are required. Such approaches have recently been applied to single-cell gene expression data^40^ and single-cell barcoding experiments^41^. However, these existing tools rely on measurements at multiple time points which are typically challenging to acquire *in vivo*. In addition, they rely on prior information regarding the global trajectories of differentiation which are aberrant in cancer, where cells can show abnormal differentiation and plasticity^42^, or do not consider variability in clone sizes^43^, a major aspect of our dataset from which we aimed to learn differential clonal behavior.

Therefore, we developed a novel dynamical modeling approach to infer the clonal differentiation behavior from the clone size distribution and their spread across clusters.. Thus, we constructed 6 concurrent models to explain the data (**Figure 5A**). These models are characterized by the cluster of origin, and directionality of the flow of cells (either uni-directed differentiation or plasticity). We characterized cells by their net proliferation (proliferation minus death) and differentiation rate. We implemented these assumptions in a dynamical model, via clone-specific proliferation-rates *β* capturing proliferation within the cluster of origin, drawn from a gamma distribution, and differentiation-rates *κ* (distinct for each direction of differentiation), drawn from mixture gamma distributions. We further assumed that proliferation also depends on the stage of differentiation (with a coefficient *r_cl_* scaling the proliferation for each cluster other than the cluster of origin) (**Figure 5B**;).

**Figure 5.**
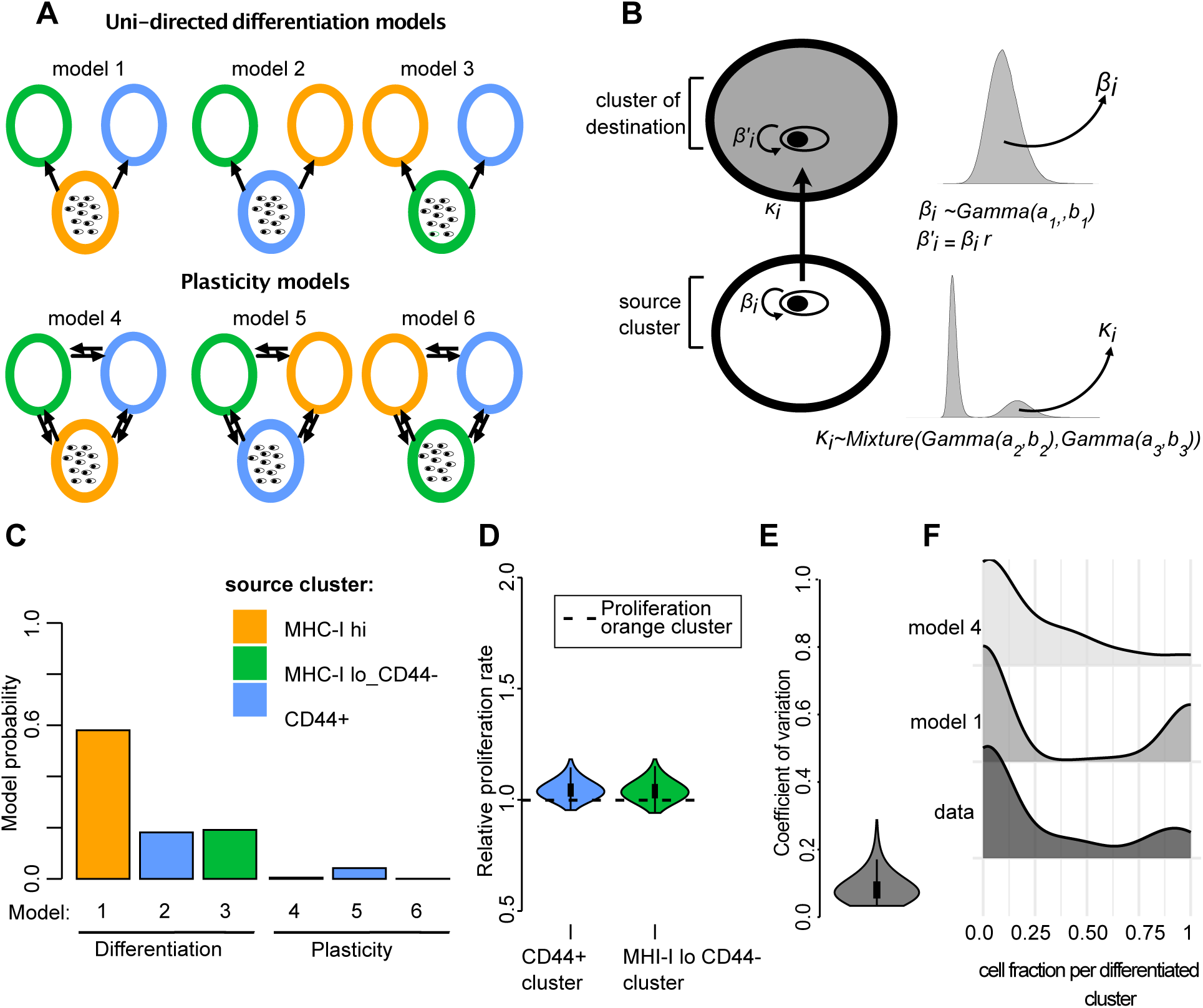
Modeling the hierarchical development in B-ALL. (A) Concurrent model of leukemia development: one-way differentiation (upper), cell plasticity (lower), columns denote different source cluster. (B) Dynamical parameters. Each clone proliferates with rate β_i_ in the source cluster and differentiates into a destination cluster with rate κ_i_, the proliferation rates are taken from a gamma distribution and the differentiation rates from a mixture gamma distribution, where some clones have a negligible differentiation into a given cluster. (C) Model probability as inferred by Approximate Bayesian Computation. (D) and (E) inferred posterior distribution for most probable model (model 1) (D), Average proliferation rates in blue and green cluster relative to the source cluster (orange); (E) coefficient of variation of the proliferation rate between clones. (F) Distribution of cell fractions per differentiated cluster as inferred for model 1, model 4 or in the data.

To simulate population growth and model observed clone size distributions, we used Approximate Bayesian Computation. A major advantage of the Bayesian approach is that the model probability and credible intervals for parameters arise directly from the distribution of the parameters corresponding to the accepted simulations (**mathematical modeling section in supplemental data**). Strikingly, the models with uni-directed “one-way” differentiation (more than 95 % probability for models 1 to 3 combined) performed far better than the models of cell plasticity (**Figure 5C**). We further found that cluster 1 was the most likely origin of differentiation (model 1), being almost 3 times as likely as the second most likely model 2. This demonstrated that the relative clone diversity between clusters is in itself sufficient to predict the directionality of the differentiation.

Notably, from the single observed time-point, we were also able to determine that proliferation of cells in clusters 2 and 3 did not differ substantially to cells in cluster 1 (**Figure 5D**). This finding was consistent with the similar experimental growth curve of the CD44^+^ and CD44^-^ cells *in vivo* (**Figure 4C and supplemental Figure 4B**), further confirming that cells differ in engraftment potential, but expand in a homogeneous manner, irrespective of the stage of differentiation. Furthermore, we could identify the coefficient of variation in proliferation between clones (ratio of the standard deviation to the mean) (**Figure 5E**). We could in particular determine the upper bound of 0.2 (95 percentile), a level of variation sufficient to model the large differences in size between clones that we observed. Of note, this rate was lower than what was previously reported for leukemic cell lines *in vitro* (0.3 - 0.41)^44^.

Finally, we investigated the reason why cell plasticity (models 4 – 6) was dismissed during the fitting procedure. For this purpose, we generated new simulations from the parameters accepted by the ABC simulations for model 1 (uni-directed differentiation) and 4 (plasticity). In doing so, we noted that the best simulations of model 4 (i.e. those that best approximating the data) systematically underestimated the variability in clone proportion for a given cluster. In contrast, such a bias was not observed for model 1 (**supplemental Figure 7**).

To better understand this bias, we visualized the distribution of clone proportions in the differentiated clusters 2 and 3 **(Figure 5F)**. This revealed that in our experimental data, the distribution was shifted towards both extremes (0 and 1), showing a bimodal distribution of cells either remaining in the source cluster or accumulate in the differentiated clusters. The same pattern was also found for the simulations of model 1. Model 4 did not replicate this concentration in the differentiated clusters: if cells can differentiate and de-differentiate in both directions (plasticity), they do not concentrate in one cluster of destination.

Taken together, our modeling approach confirmed that there is a hierarchical structure of BALL development and largely excluded the cell plasticity hypothesis.

## Discussion

Our data support the concept that B-ALL exhibits a hierarchical organization in which immature leukemic stem cells differentiate in a uni-directional way into less leukemogenic, more mature progeny. Conflicting data exist proposing both, the existence of a hierarchy with prospectively isolatable LIC populations in B-ALL^45–47^, and no hierarchical organization of the disease with high plasticity of most B-ALL cells^18,19,25,48^ with the latter becoming the predominant notion in the field.

Discriminating tumorigenic from non-tumorigenic cells based on surface markers is a fundamental principle to reconstruct the hierarchical organization of the CSC model^5^. However, reliable prospective isolation and characterization of LICs in BCP-ALL has remained challenging^4,49^. While plasticity in surface marker expression - such as for CD34, CD38^50^, CD10, CD20^51^, and CD19^52^ - has been reported, our dynamical modeling and serial transplantation experiments robustly exclude functional plasticity. Notably, we showed that while transplantation of the CD44^-^ population generated CD44^+^ cells, the phenotype of CD44^+^ cells remained stable in serial transplantation. CD44^+^ cells showed a strongly reduced efficiency in leukemia initiation in primary and secondary recipients, and about half of the mice did not demonstrate leukemic engraftment at all.

Recent work has focused on the assignment of bulk or single cell transcriptome data to specific B-cell development stages, demonstrating that ALL cases resembling early lymphocyte developmental or hematopoietic progenitor-like stages show a worse prognosis with higher likelihood of therapy resistance and relapse^53–57^. These studies hint for the existence of immature leukemic cells, however, the genealogy and trajectories of ALL cells remain unexplored by these studies. Our work reveals distinct clonal behavior, as we find that clones differ in both differentiation and proliferation properties *in vivo*, with clones being either purely self-renewing or actively differentiating, generally into one preferred differentiated population. These differentiation properties of cells were intrinsically determined, and we could identify expression markers of the differentiation bias within the immature population. Conversely, the sizes of identical clones showed no correlations between mice and proliferation was therefore likely driven by extrinsic factors such as interactions with the microenvironment and cooperation or competition between clones^58^.

Finally, our modeling framework to infer development dynamics not only confirmed the developmental hierarchy but also revealed important dynamical characteristics of the disease, notably the level of variation in proliferation between clones *in viv*o, a question which has become a focus of attention in the cancer research community^59^. This analysis demonstrates how single-cell barcoding data can inform on cancer differentiation characteristics at clonal level from a single time point.

We did not address the sources of the intrinsic differences between clones, and whether the various clonal differentiation behaviors were associated with differential epigenetic traits or mutation patterns. Furthermore, we do not propose CD44 as a general discrimination marker to enrich for distinct ALL differentiation stages, and future studies should carefully evaluate the functional and molecular properties of CD44-expressing B-ALL cells. Another open question is the generality of differentiation in B-ALL. We could find clear evidence of differentiation in our investigated patient-derived B-ALLs, however, B-ALL is a diverse disease and further studies are needed to consolidate our findings as a general feature in B-ALL.

The methodology presented here advances our understanding of B-ALL. Studies combining our experimental and analytical approach to the application of cell-based and chemotherapeutic agents may delineate the connection between cell type composition, organization, and clinical outcome leading to improved treatment strategies.

## Acknowledgements

We would like to thank Ramona Famulla for excellent technical assistance and high titer virus production. Further we thank Edyta Kowalczyk and Vadir Lopez from Becton Dickinson for their immense work with initial bioinformatic analyses and collaboration throughout the years.

This study was supported by grants from the Deutsche Jose Carreras Leukämie-Stiftung (DJCLS 11R/2020 and DJCLS 15R/2023), the Deutsche Forschungsgemeinschaft DFG (RI 2462/9-1 and RI 2462/10-1, the LOEWE Hessian Funding Program (Hessen State Ministry for Higher Education, Research and the Arts, III L5 − 519/03/03.001 – [0015] and III 5.7 - 519/03/10.001-(0004)), and the Wilhelm-Sander-Stiftung (Grant 2018-116.1). AJ was supported by a grant from the Mildred Scheel Nachwuchszentrum Frankfurt, Deutsche Krebshilfe.

Figure 4A panel has been created with BioRender.com

## Authorship Contributions

A.G., A.J., and M.A.R. planned the experiments. A.G. and T.P.S. performed the experiments under the supervision of M.A.R.. F.L., Y.M., M.K., S.G. generated and provided tools and reagents, and performed experiments. A.J. performed the analysis of the single-cell sequencing data, conceived and implemented the dynamical model under the supervision of F.B.. A.G., A.J., F.B. and M.A.R. wrote the manuscript. M.A.R. and F.B. supervised the study. All authors read and approved the manuscript.

