## supplemental information for "Differentiation hierarchy in adult B cell acute lymphoblastic leukemia at clonal resolution"

### SUPPLEMENTAL MATHEMATICAL MODEL

#### A deterministic model of clonal populations development

We seek to describe the dynamics of barcoded B-ALL cells in the bone marrow of transplanted mice. The cell population is made of distinct clones which are identified by their cell barcode.

We first estimate the total clone diversity in the right femur of each mouse using DivE[S1], a robust method based on rarefaction curves with R package version 1.3, with models 1 to 30 and default parameters. This approach yields estimates of 194 clones and 198 clones respectively for mouse 1 and mouse 2. Consequently, we model the growth of a population comprising 200 clones per mouse.

Given the very high degree of clonal diversity before transplantation with no sign of clonal dominance (supplemental Figure 1D), the number of transplanted barcoded cells (100 000), and the fact that only 200 clones were actually found to grow in the organ while thousands had been transplanted, we expect the initial number of cells per clone to be of a few units. To account for this, we draw the initial cell number of each clone from a gamma distribution with shape 10 and scale 0.5 (mean 5 cells and variance 2.5 cells).

The three clusters observed after transplantation are in particular characterized by differential surface expression of HLA-ABC molecules. Flow cytometry measurements of the cells in vitro showed no variable expression of HLA-ABC (supplemental Figure 2C), we consequently choose to initialize the simulation with all the cells contained within a single cluster with the most probable cluster of origin being determined by model selection.

Successive measurements of percentage of leukemic cells in the blood during leukemogenesis (Figure 4C and supplemental Figure 5B) demonstrate an exponential growth of the leukemic cell population, we therefore consider that each clone $i$ grows exponentially and that the system can be modeled by a system of ordinary differential equations. At a given time, each cluster $j$ (with $1 \leq j \leq 3$) is made of $n$ clones $i$ of size $C_{j,i}$.

The proliferation rates may vary between clones and the cluster may also have an influence on proliferation rates. To represent this variability, the proliferation parameters per clone in the source cluster $\beta_{1,i}$ are drawn from gamma distributions where the shape parameter $\alpha_\beta$ determines the variability of the proliferation rate between clones and the relative scale parameter $\theta_\beta$, with mean proliferation rate per clone $\mu_\beta = \theta_\beta/\alpha_\beta$.

For the differentiated clusters 2 and 3, the proliferation rate is then affected by a scaling factor $r_j$ so that the proliferation rate in a differentiated cluster $\beta_{j,i} = r_j \cdot \beta_{1,i}$.

We definefor each clone $i$ the rates $\kappa_{1,i}$, $\kappa_{2,i}$ and $\kappa_{3,i}$ ,respectively the differentiation rates from cluster 1 (source cluster) to cluster 2, cluster 1 to cluster 3, and cluster 2 to cluster 3. We observed from the repartition of cells between cluster (Figure 2C) that clones are highly concentrated into one cluster so we can reasonably assume that many clones do not differentiate, or will often exclusively differentiate into one destination cluster. To model this phenomenon, we draw the differentiation rates $\kappa_{1,i}$,$\kappa_{2,i}$ and $\kappa_{3,i}$ from mixture distributions with clones having alternatively an active differentiation rate taken from one distribution or a background differentiation rate (Figure 5B) with $p_1$, $p_2$ and $p_3$ the probabilities to draw from the background distribution for each $\kappa$. The gamma distribution of background differentiation rates is shared for every $\kappa$ and has for parameters $\alpha_\kappa$ and $\theta_{\kappa,0}$ with mean rate $\mu_{\kappa,0}$; the 3 distributions of active differentiation rates have for parameters $\alpha_\kappa$ and respectively $\theta_{\kappa_1}$,$\theta_{\kappa_2}$ and $\theta_{\kappa_3}$.

The parameters of these distributions of proliferation and differentiation rates are estimated from the data for the models described below.

*One-way differentiation model*

We first model, a system where cells starting in a cluster 1 can proliferate and differentiate uni-directionally from cluster 1 into clusters 2 and 3. We describe the dynamics of each clone as:

$$\begin{aligned}\frac{d}{dt}C_{1,i,t} &= C_{1,i,t} \cdot (\beta_{1,i} - \kappa_{1,i} - \kappa_{2,i}) \\ \frac{d}{dt}C_{2,i,t} &= C_{2,i,t} \cdot \beta_{2,i} + C_{1,i,t} \cdot \kappa_{1,i} \\ \frac{d}{dt}C_{3,i,t} &= C_{3,i,t} \cdot \beta_{3,i} + C_{1,i,t} \cdot \kappa_{2,i}\end{aligned} \quad (1)$$

This linear system has an analytical solution given by:

$$\begin{aligned}C_{1,i,t} &= C_{1,i,0} \cdot \exp\left[-t \cdot (\kappa_{1,i} + \kappa_{2,i} - \beta_{1,i})\right] \\ C_{2,i,t} &= \frac{\kappa_{1,i} \cdot C_{1,i,0} \cdot \exp(\beta_{2,i} \cdot t)}{\kappa_{1,i} + \kappa_{2,i} - \beta_{1,i} + \beta_{2,i}} - \frac{\kappa_{1,i} \cdot C_{1,i,0} \cdot \exp\left[-t \cdot (\kappa_{1,i} + \kappa_{2,i} - \beta_{1,i})\right]}{\kappa_{1,i} + \kappa_{2,i} - \beta_{1,i} + \beta_{2,i}} \\ C_{3,i,t} &= \frac{\kappa_{2,i} \cdot C_{1,i,0} \cdot \exp(\beta_{3,i} \cdot t)}{\kappa_{1,i} + \kappa_{2,i} - \beta_{1,i} + \beta_{3,i}} - \frac{\kappa_{2,i} \cdot C_{1,i,0} \cdot \exp\left[-t \cdot (\kappa_{1,i} + \kappa_{2,i} - \beta_{1,i})\right]}{\kappa_{1,i} + \kappa_{2,i} - \beta_{1,i} + \beta_{3,i}}\end{aligned} \quad (2)$$

*Plasticity model*

We also introduce another model of differentiation including plasticity, where differentiation is reversible and the differentiations rates work in both directions. We also allow transdifferentiation between cluster 2 and 3 with rate $\kappa_{3,i}$.

$$\begin{aligned}\frac{d}{dt}C_{1,i,t} &= C_{1,i,t} \cdot (\beta_{1,i} - \kappa_{1,i} - \kappa_{2,i}) + C_{2,i,t} \cdot \kappa_{1,i} + C_{3,i,t} \cdot \kappa_{2,i} \\ \frac{d}{dt}C_{2,i,t} &= C_{2,i,t} \cdot (\beta_{2,i} - \kappa_{1,i} - \kappa_{3,i}) + C_{1,i,t} \cdot \kappa_{1,i} + C_{3,i,t} \cdot \kappa_{3,i} \\ \frac{d}{dt}C_{3,i,t} &= C_{3,i,t} \cdot (\beta_{3,i} - \kappa_{2,i} - \kappa_{3,i}) + C_{1,i,t} \cdot \kappa_{2,i} + C_{2,i,t} \cdot \kappa_{3,i}\end{aligned} \quad (3)$$

This system has no practical analytical solution.

**Parameter estimation**

We compute the posterior distribution for parameters using the Sequential Monte Carlo Approximate Bayesian Computation algorithm implemented by the package pyABC[S2] version 0.12.15 with the following parameters:

- population size = 8000
- fitting stops when an acceptance rate of 0.01 is reached
- standard acceptance threshold scheduling
- MulticoreEvalParallel sampler

*simulating the models*

We generate simulations for 6 models, that is, for each of the two above mentioned models, we simulate three instances, with either the MHC-I, the CD44+ or the MHC-I lo CD44- neg cluster considered as source cluster.

For each of the two mice replicates, we generate model predictions at day 50 following cell transplantation as in the experimental dataset. For the One-way differentiation model, we compute directly the solutions for each clone using the analytical solution. For the plasticity model, we generate numerical solutions of the system of ordinary differential equations using the ivp solver of python package scipy version 1.14.1.

*summary statistics*

From the cell counts per clone at day 50, we derive summary statistics to compare the model prediction to the data.

For each mouse, we first restrict the data to 42 largest clones to avoid overemphasis on small clones which are most subject to sampling error. From Figure 3A, we know there are characteristic patterns of clone distribution across cluster so we expect the counts to be interdependent. Consequently, we use the proportion of cells per cluster for each clone rather than counts $c_{ij} = C_{ij}/C_i$, with $C_i$ the total number of cells of clone $i$ in the dataset and $C_{ij}$ the number of cells of clone $i$ in cluster $j$.

We also compute the relative size of each clone $c_i$ compared to the whole dataset $N$.

$$c_i = \frac{C_i}{N}$$

We order clones by decreasing size:

$$c_1 \geq c_2 \geq \cdots \geq c_n$$

The repartition of a clone between clusters at a specific rank is likely due to chance, we summarize the information by grouping the ordered clones into $m$ bins of 6 clones each.

For each bin $k$, we compute the average cell fraction per cluster $\bar{c}_{k,j}$ and the average proportion of cells per clone $\bar{c}_k$ .

$$\bar{c}_{k,j} = \frac{1}{6}\sum_{k=1}^{m} c_{k,j} :$$

and the relative average relative size of the clones in bin $k$:

$$\bar{c}_k = \frac{1}{6}\sum_{k=1}^{m} c_k$$

Similarly compute the standard deviations $\sigma_{k,j}$ and $\sigma_k$.

Given $\bar{c}_k^{pred}$, $\bar{c}_{k,j}^{pred}$, $\sigma_k^{pred}$ and $\sigma_{k,j}^{pred}$ which are calculated in the same way from the model simulations described in the above section, we compute the following distance to determine the posterior distributions:

$$distance = \sum_{k=1}^{b} \mid \bar{c}_k^{pred} - \bar{c}_k \mid + \sum_{j=1}^{3}\sum_{k=1}^{b} \mid \bar{c}_{k,j}^{pred} - \bar{c}_{k,j} \mid + \sum_{k=1}^{b} \mid \sigma_k^{pred} - \sigma_k \mid + \sum_{j=1}^{3}\sum_{k=1}^{b} \mid \sigma_{k,j}^{pred} - \sigma_{k,j} \mid$$

### Model parameters and priors

With little initial information on the parameters range in our particular system, we generally choose uniform prior distributions with reasonable lower and upper bounds for a proliferating cell system. Of note, Given that cells are generally concentrated into one cluster, we assume that at least 50% of the clones are mostly self-renewing will not be able to differentiate actively. Furthermore, following the observation that every cluster is highly proliferative (supplemental Figure 2F), we suspect the proliferation will not vary dramatically between clusters and therefore, we set a normal prior distribution for $r_2$ and $r_3$ centered on 1, i.e the proliferation in the source cluster. All parameters and priors are presented in supplemental Table 2.

### Credible interval for parameters as inferred by ABC for model 1

from the above mentioned parameters, we derive $1/\sqrt{\alpha_\beta}$, the coefficient of variation of proliferation rates and $1/\sqrt{\alpha_\kappa}$, the coefficient of variation of the differentiation rates.The inferred credible intervals for parameters are presented in supplemental Table 3.

### SUPPLEMENTAL METHODS

#### Generation of the expression barcoding vector using the 3Cs technology

The expression barcoding vector was generated with the cloning-free 3Cs technology as described by Wegner et al.[S3] The lentiviral vector pSEW[S4] served as the backbone of the expression barcoding vector. First, the f1 origin of replication was introduced into the backbone for the 3Cs cloning-free technology. Then, the open reading frame of mCherry followed by the insertion site of the barcode sequence and a PolyA signal at the 3´untranslated region was cloned behind a bidirectional human PGK promoter driving reporter and barcode expression in antisense. The barcodes were designed according to the ClonTracer library[S5], containing a 30mer a semi-random order of weak (W) and strong (S) base-pairing.

Fifty nanograms of the vector were transformed into the E. coli strain CJ236. Transformants were grown overnight at 37 °C on LB agar plates containing chloramphenicol (25 µg/ml) and ampicillin (100 µg/ml). Four single colonies were picked and cultured in 1 ml of 2xYT medium supplemented with ampicillin (100 µg/ml) and 1 × $10^8$ pfu of M13KO7 helper phage (New England Biolabs, N0315). After 3 h of incubation at 200 rpm and 37 °C, kanamycin (50 µg/ml) was added to select for M13KO7-infected bacteria. Cultures were incubated for an additional 5 h under the same conditions and then transferred to 30 ml of 2xYT medium with ampicillin (100 µg/ml) and kanamycin (50 µg/ml). After 20 h of growth at 200 rpm and 37 °C, the culture was centrifuged in a Beckman JA-12 rotor for 10 min at 12,000 g and 4 °C. The supernatant was mixed with 6 ml PEG/NaCl buffer (20% PEG 8000, 2.5 M NaCl) and incubated for 60 min at room temperature. The phage suspension was centrifuged for 10 min at 12,000 g and 4 °C, and the pellet was resuspended in 1 ml PBS (Sigma-Aldrich, D8662). After centrifugation for 5 min at 16,000 g, the phage-containing supernatant was collected and stored at 4 °C until ssDNA purification. Circular ssDNA was isolated using the E.Z.N.A. M13 DNA Mini Kit (Omega Bio-Tek, D69001-01) following the manufacturer's protocol.

The semi-random barcode-containing oligonucleotides (5´-ACTGACTGCAGTCTGAGTCTGACAG WSWSWSWSWSWSWSWSWSWSWSWSWSWSWSAGCAGAGCTACGCACTCTATGCTAG-3´) were ordered from Sigma Aldrich. To phosphorylate and anneal the oligonucleotides, 154 ng of randomized oligonucleotide was phosphorylated in a 20 µl reaction containing 2 µl 10× TM Buffer (0.1 M $MgCl_2$, 0.5 M Tris–HCl, pH 7.5), 2 µl 10 mM ATP (New England Biolabs, 756), 1 µl 100 mM DTT (Cell Signaling Technology Europe, 7016), and 2 µl 10,000 U/ml T4 Polynucleotide Kinase (New England Biolabs, M0201). The mixture was incubated at 37 °C for 1 h. Subsequently, 20 µg of ssDNA was combined with 25 µl 10× TM buffer, 20 µl of phosphorylated oligonucleotide, and water to a final volume of 250 µl. This mixture was denatured at 98 °C for 1 min, cooled gradually (0.1 °C/s) to 45 °C for 5 min, and then further cooled to room temperature for 10 min.

To the annealed ssDNA–oligonucleotide mixture, 10 µl 10 mM ATP, 10 µl 100 mM dNTP mix (Carl Roth GmbH, 0178.1/2), 15 µl 100 mM DTT, 2000 U T4 DNA ligase (New England Biolabs, M0202), and 30 U T7 DNA polymerase (New England Biolabs, M0274) were added. The reaction was incubated overnight at RT. The 3Cs product was purified with the DNA Clean & Concentrator-25 kit (Zymo Research, D4034) according to the manufacturer's instructions. The purified product was electroporated into NEB 10-beta Electrocompetent E. coli (C3020K) and cultured overnight in 250 ml LB medium with ampicillin at 37 °C for 16–18 h. Plasmid DNA was subsequently isolated using the QIAGEN Plasmid Plus Maxi Kit.

**Bulk RNA Sequencing**

Isolation of cellular RNA from a minimum of 10,000 sorted cells was performed using the RNeasy Micro Kit (Qiagen, Hilden, Germany) according to the manufacturer's protocol. Afterwards, 50 ng total RNA was used as the starting material using the SMARTer Universal Low Input RNA Kit for Sequencing (Takara Bio, Kusatsu, Japan) according to the manufacturer's protocol. Sequencing of final libraries was performed using a NextSeq2000 Sequencer (Illumina, San Diego, CA, USA) with 72 bp single-end settings.

Acquired raw reads were evaluated for quality, adapter content, and duplication rates using FASTQC v0.11.9. Only reads longer than 15 nucleotides were preserved for subsequent analyses, mapped to the Ensembl human genome version hg38 using STAR v2.7.10a. The filtered raw count matrix was normalized with DESeq2 v1.36.0. Finally, genes with an average count > 5, multiple testing adjusted p-value < 0.05, and a log2FC between -0.585 and +0.585 were selected as significantly differentially expressed.

**Single-cell Transcriptome and Epitope Sequencing**

Before sorting, cells from different animals were incubated with Sample Tags for multiplexing using the Hu-Single Cell Sample Multiplexing Kit (BD, Franklin Lakes, NJ, USA). After sorting human $CD45^+$, $mCherry^+$ cells, samples were stained with 49 AbSeq oligo-conjugated antibodies (**Table S1**, BD, Franklin Lakes, NJ, USA) for single-cell transcriptome and epitope sequencing. Subsequently, cells were stained with Draq7 (BD, Franklin Lakes, NJ, USA) and Calcein AM (2 mM, BD, Franklin Lakes, NJ, USA) to assess their viability, before 40,000 cells were loaded onto one Rhapsody microwell cartridge (BD, Franklin Lakes, NJ, USA) and processed according to the manufacturer's protocol. After retrieving beads from the cartridge, the cDNA was generated and WTA, Sample Tag and AbSeq libraries were prepared using the Rhapsody WTA Amplification kit (BD, Franklin Lakes, NJ, USA). The ClonTracer barcode information of single cells was obtained by preparing a library using the Rhapsody Targeted mRNA kit with custom primers binding to sequences flanking the barcode (BD, Franklin Lakes, NJ, USA). Final libraries were sequenced with 20 % PhiX in multiple runs (75 bp paired-end) using a NextSeq2000 sequencer (Illumina, San Diego, CA, USA).

FASTQ sequencing and reference files were uploaded to the SevenBridges platform (Seven Bridges Genomics, Boston, MA, USA, https://www.sevenbridges.com). The Rhapsody WTA analysis pipeline was then used with default settings to process the data. In brief, low-quality read pairs were excluded, prior to mapping the remaining R1 reads to identify the cell label and unique molecular identifier (UMI) sequence. Subsequently, R2 reads were aligned to reference sequences (GRCh38.p12), and reads sharing identical cell labels, UMI sequences, and reference genes were collapsed into single molecules. A recursive substitution error correction (RSEC) algorithm was used to correct sequencing errors in molecule counts. Following this, Sample Tags were assigned to cells, and cell counts were calculated, generating RSEC-adjusted molecule count matrices.

**Single-cell data analysis**

The datasets were loaded in R as SingleCellExperiment objects. Low quality cells were excluded based on feature number per cell, UMI counts per cell using perCellQCFilters() from the package scran v1.30.2 with default parameters and cells expressing more than 25% of mitochondrial genes were further excluded. Single-cell RNA seq data was normalised by deconvolution using computeSumFactors() and logNormCounts() from the package scran.

The AbsSeq data was normalized separately. Cells with no AbSeq expression where also discarded. The background signal in the AbSeq data was determined using ambientProfileBimodal() from the package DropletUtils v 1.22.0, the size factor for each cell was estimated with the function medianSizeFactors() taking into account the background noise calculated above. Cells with a median size factor equal to 0 were further excluded at this stage. The counts were then log2 normalized using NormLogCounts().

We performed an initial clustering to use for doublet calling, for this purpose, we performed PCA on transcriptome and AbSeq data and performed multiUMAP combining both layers using 6 principal components for each layer. We then clustered the data using the MultiUMAP with Louvain clustering at resolution 0.03 for BV and 0.01 for each of the 2 CR samples respectively.

We then called doublets using the main function of scDblFinder v.1.16.0. We used the expected percentage of doublets as provided in the report of the BD Rhapsody system, respectively (4.4% and 5.4% for the patient CR-derived samples and 5% for the patient BV-derived B-ALL sample) and also provided the cluster information to the algorithm.

For integration of the two patient CR-derived samples, the samples were rescaled using rescaleBatches() from the package Batchelor v 1.18.1, PCA and UMAP revealed no further batch effect to be corrected.

For the final visualization of the single-cell data, we removed the cell-cycle effect from the transcriptome using Seurat v5.1.0. The datasets were converted into SeuratObject, Cell cycles scores were calculated based on S and G2M signature genes defined in the Seurat package. A PCA was computed using Seurat cell-cycle corrected and scaled values and a MultiUMAP was computed as mentioned above. The final clustering was then performed as above. We then performed diffusion map on the cell cycle corrected transcriptome with the package destiny v 3.16.0 with the parameters PCs=4 and k=90.

For the analyses requiring python packages (PAGA), the dataset was converted into .h5ad format with the package ZellKonverter v 1.12.1. The dataset was then opened in python v3.10 as anndata and processed with Scanpy v 1.10.3.

For PAGA analysis, the neighborhood graph was computed with 4 Principal Components a connectivity between cluster was then inferred and cluster to cluster connectivity higher than 0.1 was plotted.

**Assignment to B cell development stage**

The SingleCellExperiments were converted to Seurat objects as mentioned above and  cells were projected on the B-ALL development map as described in https://github.com/andygxzeng/b_development_map.

The stages Pre-Pro-B and Pre-Pro-B Cycling` and Pro-B VDJ`, `Pro-B Cycling` were merged as Pre-Pro-B and Pro-B respectively. And were then computed the proportion of cells in each stage of B cell development (“Pre-Pro-B”, “Pro-B”, “Large Pre-B”, “Small Pre-B”, “Immature B”, “Mature B”).

**Lineage Barcode extraction and filtering**

Lineage barcodes were extracted from the BAM files the reads which had been mapped to the “user_seq_masked_barcode” feature using Samtools. Barcode filtering was then performed in R with the packages Rsamtools v. 2.18.0 and DECIPHER v 2.30.0 (filtering pipeline available:

https://github.com/AdrienJolly/B_ALL_differentiation). (i) Reads containing adapter sequence "TGTACAACTAGGCGGCCGCTAGACTGACTGCAGTCTGAGTCTGACAG" were trimmed using function TrimDNA with default settings and reads of size 26 to 28 bases following successful trimming were conserved. (ii) The 26 bases directly following the adapter sequences were used as lineage barcodes; (iii) in order to exclude cells carrying multiple barcodes ( ambient RNA contamination), for each cells, a consensus barcode sequence was defined using the function ConsensusSequence() and only cells with a consensus sequence representing at least 90% of all barcode reads of the cell were considered barcoded; (iv) cells with barcodes supported by less than 4 reads in the dataset were not considered bacoded; (v) finally, barcodes present in only one cell were then collapsed with a larger clones (barcode present in more than one cell) if : (1.) They were supported by less than 4 UMIs and, (2.) They had a Hamming distance of 1 or 2 with the larger barcode.

**Learning predictors of differentiation**

As shown in **Figures 2E** and **Fig S2F**, we grouped clones of size 20 and more into classes based on their distribution across clusters, The classes were either determined based on the two first principal components (PCA performed with FactoMineR R package version 2.11; **Figure 2E**) or based on hierarchical clustering with Euclidean distance (R package pheatmap version 1.0.12, **Figure S2F)**

We used XGBoost R package v 1.7.8.1 to learn predictors of the differentiation classes from their transcriptome in the source cluster. For this purpose, we selected, for each patient-derived B-ALL, cells of the 3 (for CR,) or 2 (for BV) classes in cluster 1, and split them into 90% training set and 10% test set. For each patient-derived B-ALL the models were trained using the top 500 highly variable genes determined from the above-mentioned cells of interest.

The datasets are characterized by high class imbalance so we to generate subsamples from the training set to get a representative sample of comparable size for each class.

In order to get similar levels of transcriptomic variability per class, and given the expected high correlation between cells of a given clone[26], we selected a certain number of cells from the smallest class and then determined the number of cells of the other classes based on the relative clonal diversity of each class. Namely, we computed Shannon's entropy for each class and used the ratio of this entropy to that of the smallest class as a scaling factor to determine the number of cells to be used for training.

We iteratively split the global training set into training with the number of cells as described above, and validation sets randomly 50 times. At each iteration, XGboost was run on the log2 normalized counts with the following parameters: max.depth = 100, eta = 1, nrounds = 100, objective = "multi:softmax". Finally, we selected the best of the 50 models based on average recall for the differentiating classes (classes overrepresented in the mature clusters).

In order to get the statistical significance of the prediction, we then iteratively permuted the classes within the global training set and repeated the whole training procedure as described above 1000 times. The balanced accuracy obtained from the dataset was compared to the distribution obtained from the permutations to determine the statistical significance of the true model.

We selected the 20 best predictors based on prediction gain and computed the relative expression for the cells of the different classes in cluster 1 using scaleData from the Seurat package as above, and then averaged this relative expression per class (**Figures 2G and S2G**).

**Statistical analysis**

Statistical analyses were performed in GraphPad Prism (version 10.4.1), R (version 4.3.0) and Python (version 3.10). All sample data was tested for normality of distribution to determine the appropriate test.

For **Figure 2D**, in order to determine the statistical significance, the clones shared by the two mice with more than 10 cells per mouse were selected, and the clone labels were permuted with replacement 1000 times. At each permutation the sum of the Euclidean distances per clone between cluster distributions was computed. It was finally compared to the true distance between mice to obtain the p-value.

The statistical test performed on the XGboost models is detailed in "Learning predictors of differentiation" section above.

The employed tests, value of n, multiple comparison corrections, and significance levels are otherwise specified and shown in the figure legends. "SEM" stands for Standard Error of the Mean, "SD" for Standard Deviation.

| Antigen | Clone | Cat# |
|---|---|---|
| CD1a | HI149 | 940063 |
| CD1c | F10/21A3 | 940083 |
| CD3 | SK7 | 940000 |
| CD7 | M-T701 | 940029 |
| CD9 | M-L13 | 940078 |
| CD10 | HI10A | 940045 |
| CD11b | M1/70 | 940008 |
| CD11c | B-LY6 | 940024 |
| CD13 | WM15 | 940044 |
| CD15 | W6D3 | 940274 |
| CD14 | MPHIP9 | 940005 |
| CD19 | SJ25C1 | 940004 |
| CD22 | HIB22 | 940273 |
| CD25 | 2A3 | 940009 |
| CD26 | M-A261 | 940101 |
| CD32 | FLI8.26 | 940069 |
| CD33 | WM53 | 940031 |
| CD34 | 581 | 940021 |
| CD38 | HIT2 | 940013 |
| CD44 | L178 | 940251 |
| CD45 | HI30 | 940002 |
| CD45RA | HI100 | 940011 |
| CD45RO | UCHL1 | 940022 |
| CD47 | B6H12 | 940082 |
| CD56 | NCAM16.2 | 940007 |
| CD62L | DREG-56 | 940041 |
| CD64 | MD22 | 940262 |
| CD81 | JS-81 | 940052 |
| CD90 | 5E10 | 940032 |
| CD93 | R139 | 940215 |
| CD96 | 6F9 | 940272 |
| CD117 (c-kit) | YB5.B8 | 940051 |
| CD123 (IL3Ra) | 7G3 | 940020 |
| CD124 | HIL4R-M57 | 940092 |
| CD126 (IL6R) | M5 | 940090 |
| CD133 | W6B3C1 | 940095 |
| CD137 | 4B4-1 | 940055 |
| CD155 (PVR) | TX24 | 940102 |
| CD184 (CXCR4) | 12G5 | 940056 |
| CD235a/b | GA-R2 (HIR2) | 940040 |
| CD273 (PDL2) | MIH18 | 940071 |
| CD274 (B7-H1) | MIH1 | 940035 |
| HLA-DR | G46-6 | 940010 |
| HLA-A,B,C | G46-2.6 | 940062 |
| CD366 (TIM-3) | 7D3 | 940066 |
| B7-H4 | MIH43 | 940100 |
| CD371 (CLL-1) | 50C1 | 940212 |
| MTO - MICA/B | 6D4 | Custom 460032 |
| GPR56 | CG4 | Custom 460005 |

**Supplemental Table 1. AbSeq Panel**

| Parameter | Description | Prior distribution |
|---|---|---|
| $\mu_{\beta_1}$ | mean proliferation rate ($d^{-1}$) in cluster 1 | $Uniform(0.1, 0.7)$ |
| $\alpha_{\beta_1}$ | shape parameter for the gamma distrib. of prol. rates $\beta_1$ | $Uniform(5, 1000)$ |
| $r_2$ | proliferation scaling factor for cluster 2 | $N(1, 0.3)$ |
| $r_3$ | proliferation scaling factor for cluster 3 | $N(1, 0.3)$ |
| $\mu_{\kappa,0}$ | mean background differentiation rate ($d^{-1}$) | $Uniform(10^{-7}, 10^{-6})$ |
| $\alpha_\kappa$ | shape parameter for the gamma distributions of diff. rates | $Uniform(200, 600)$ |
| $\mu_{\kappa_1}$ | mean active differentiation rate ($d^{-1}$) between clusters 1 and 2 | $Uniform(0.1, 0.7)$ |
| $\mu_{\kappa_2}$ | mean active differentiation rate ($d^{-1}$) between clusters 1 and 3 | $Uniform(0.1, 0.7)$ |
| $\mu_{\kappa_3}$ | mean active differentiation rate ($d^{-1}$) between clusters 2 and 3 | $Uniform(0.1, 0.7)$ |
| $p_1$ | probability to sample from background differentiation distrib. for parameters $\kappa_1$ | $Uniform(0.5, 0.9)$ |
| $p_2$ | probability to sample from background differentiation distrib. for parameters $\kappa_2$ | $Uniform(0.5, 0.9)$ |
| $p_3$ | probability to sample from background differentiation distrib. for parameters $\kappa_3$ | $Uniform(0.5, 1)$ |

**Supplemental Table 2. Model parameters and their prior distributions**

| Parameter | median + credible interval |
|---|---|
| $\mu_\beta$ | $0.43(0.18, 0.65)$ |
| $1/\sqrt{\alpha_\beta}$ | $0.07(0.04, 0.17)$ |
| $r_2$ | $1.04(0.99, 1.12)$ |
| $r_3$ | $1.04(0.97, 1.12)$ |
| $\mu_{\kappa,0}$ | $1.5 \cdot 10^{-6}(1.1 \cdot 10^{-6}, 1.9 \cdot 10^{-6})$ |
| $1/\sqrt{\alpha_\kappa}$ | $0.04(0.036, 0.06)$ |
| $\mu_{\kappa_1}$ | $0.39(0.15, 0.65)$ |
| $\mu_{\kappa_2}$ | $0.39(0.15, 0.65)$ |
| $\mu_{\kappa_3}$ | $0.4(0.14, 0.65)$ |
| $p_1$ | $0.72(0.54, 0.86)$ |
| $p_2$ | $0.78(0.57, 0.89)$ |
| $p_3$ | $0.75(0.54, 0.96)$ |

**Supplemental Table 3. Model parameters medians and 90% credible interval**

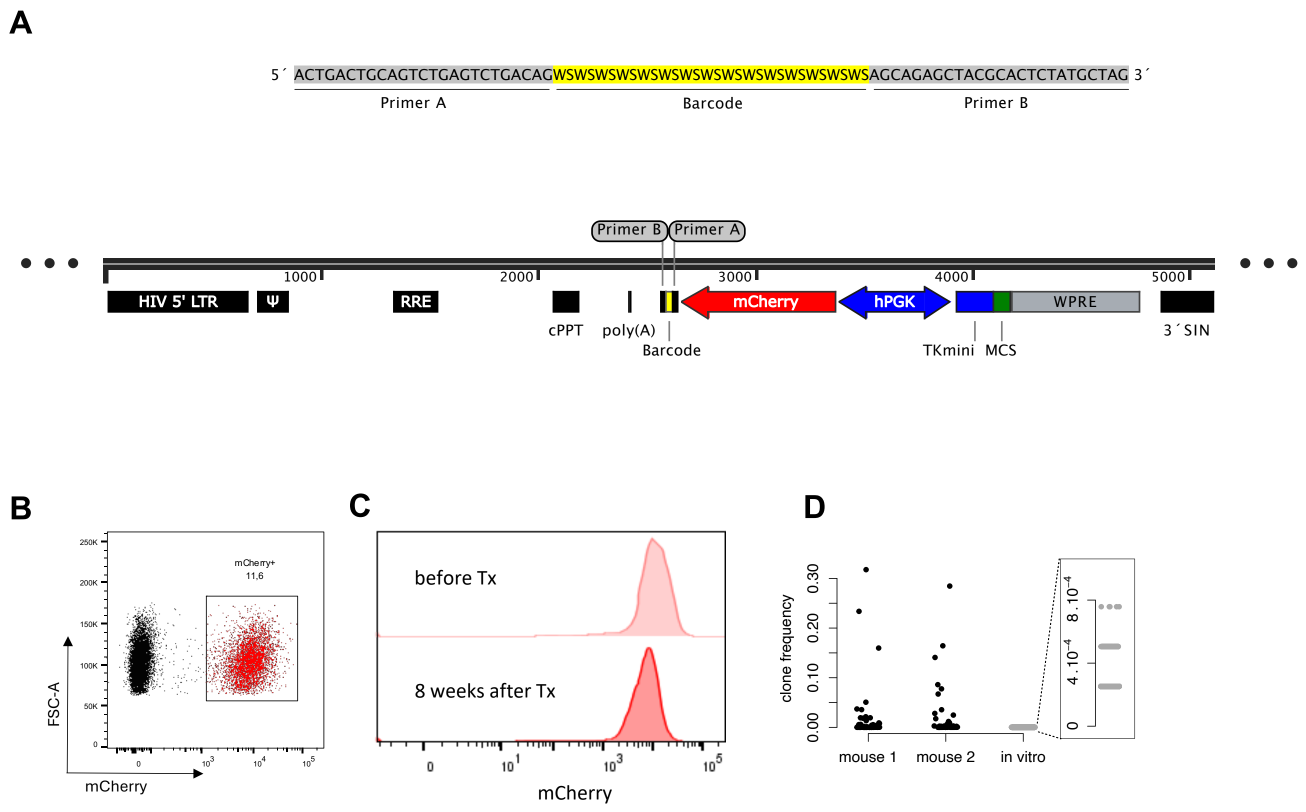


**Supplemental Figure 1. Expression barcoding vector**

**(A)** Schematic representation of the expression barcoding vector. Only the lentiviral integration part is displayed here. The 30mer barcode, flanked by two primer sites (Primer A & Primer B), was inserted upstream of the poly(A) signal into the 3' untranslated region (UTR) after the mCherry open reading frame. (**B**) Transduction of patient CR-derived B-ALL cells with the barcoding construct resulted in strong and robust mCherry expression. (**C**) Representative histogram of mCherry fluorescence intensity before and 8 weeks after transplantation of barcoded cells into NSG mice. (**D**) Clone frequency observed after leukemogenesis in mouse 1 and mouse 2 and before transplantation (*in vitro*) with zoom on low frequency clones for *in vitro* cultivated sample. HIV, human immunodeficiency virus; LTR, long terminal repeat; RRE, Rev response element; cPPT, central polypurine tract; hPGK, human phosphoglycerate kinase 1 promoter; TKmini, Thymidine kinase mini promoter; MCS, multiple cloning site; WPRE, Woodchuck Hepatitis Virus Posttranscriptional Regulatory Element; SIN, self-inactivating;

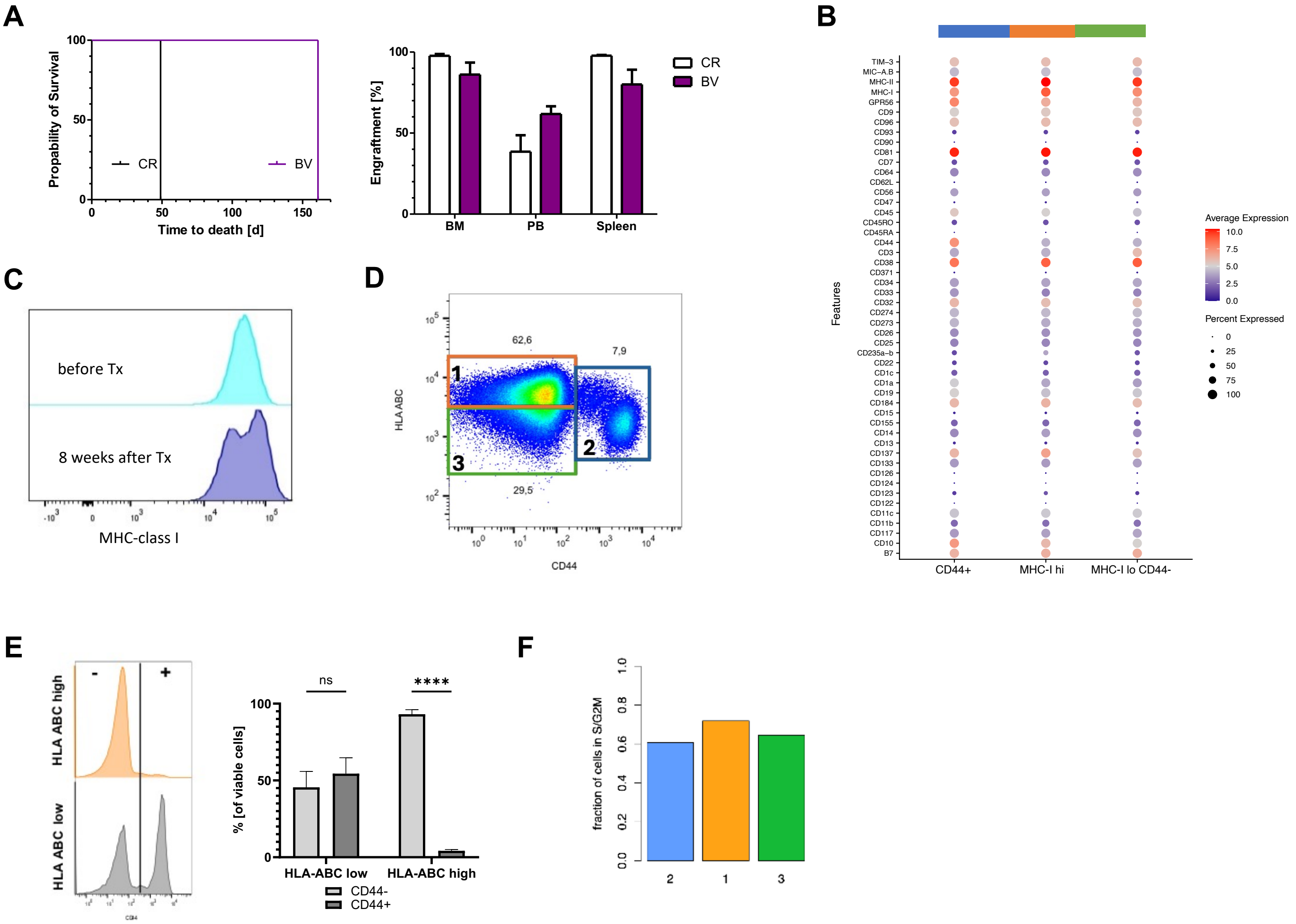


**Supplemental Figure 2. Engraftment and molecular features of barcoded B-ALL cells. Related to Figure 1.**

(**A**) Kaplan-Meier-Survival analysis of mice transplanted with patient CR (black) or BV (violet) derived B-ALLs. Engraftment of CR (white) and BV (violet) cells in Bone Marrow (BM), peripheral blood (PB) and spleen at endanalysis (n = 4-5 mice). **(B)** Normalized AbSeq expression across clusters for patient CR-derived cells. (**C**) Representative Histogram of MHC-class I fluorescence intensity before and 8 weeks after transplantation of barcoded cells into NSG mice. (**D**) Flow cytometry-based analysis of CD44 and HLA-ABC surface expression of patient CR-derived cells after transplantation. In the representative dot plot, three clusters are identified. (**E**) Representative Histogram of CD44 expression of HLA-ABC high/low patient CR-derived cells. (**F**) Fraction of cells from the three identified clusters in S/G2M cell cycle phase as inferred by Seurat.

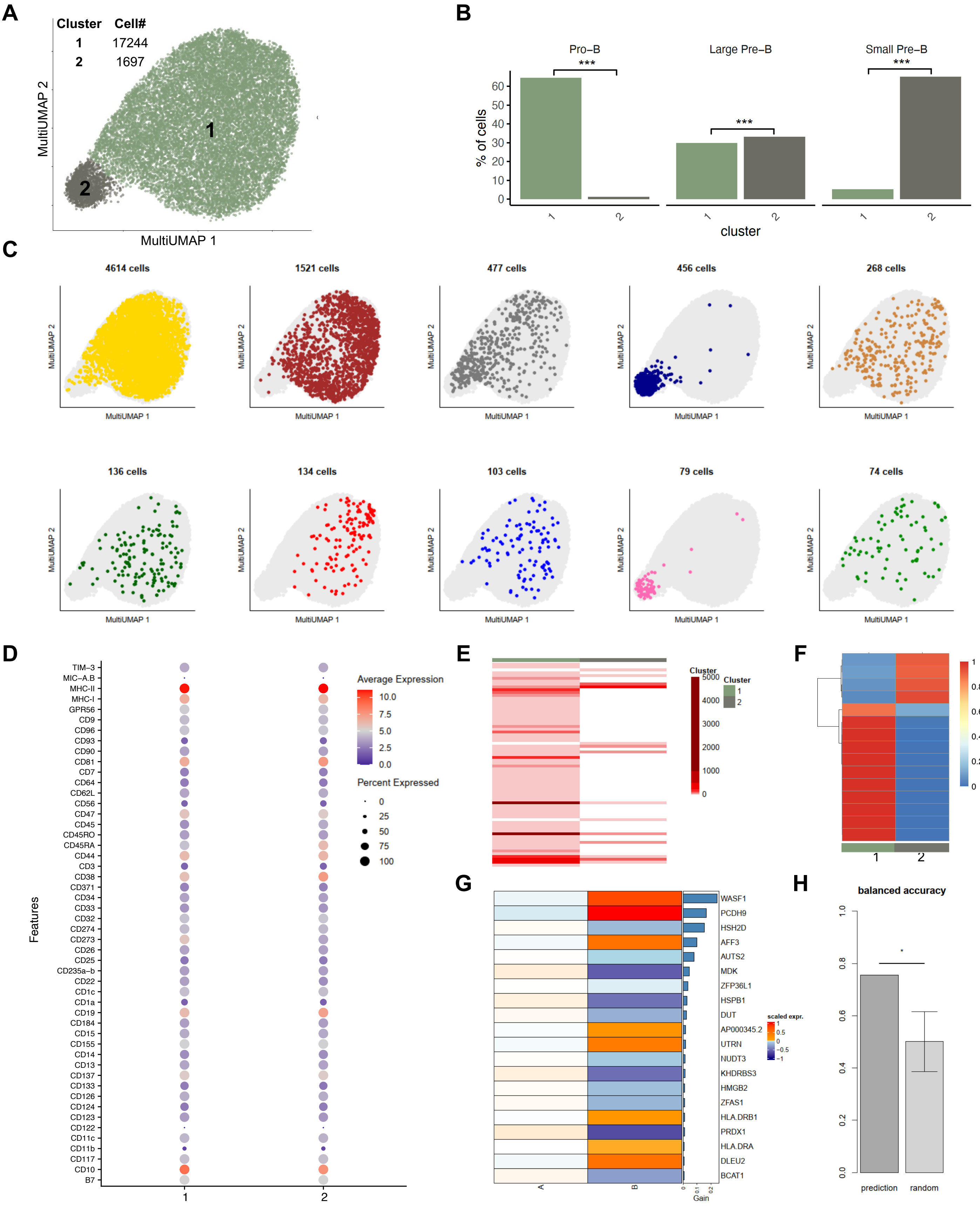
A
Cluster Cell#
1 17244
2 1697
MultiUMAP 2
MultiUMAP 1
B
Pro–B
Large Pre–B
Small Pre–B
***
% of cells
cluster
C
4614 cells
1521 cells
477 cells
456 cells
268 cells
136 cells
134 cells
103 cells
79 cells
74 cells
D
Features
Average Expression
Percent Expressed
TIM–3
MIC–A.B
MHC-II
MHC-I
GPR56
CD9
CD96
CD93
CD90
CD81
CD7
CD64
CD62L
CD56
CD47
CD45
CD45RO
CD45RA
CD44
CD3
CD38
CD371
CD34
CD33
CD32
CD274
CD273
CD26
CD25
CD235a–b
CD22
CD1c
CD1a
CD19
CD184
CD15
CD155
CD14
CD13
CD137
CD133
CD126
CD124
CD123
CD122
CD11c
CD11b
CD117
CD10
B7
E
Cluster
F
G
WASF1
PCDH9
HSH2D
AFF3
AUTS2
MDK
ZFP36L1
HSPB1
DUT
AP000345.2
UTRN
NUDT3
KHDRBS3
HMGB2
ZFAS1
HLA.DRB1
PRDX1
HLA.DRA
DLEU2
BCAT1
scaled expr.
Gain
H
balanced accuracy
prediction
random

**Supplemental Figure 3. Analysis of CITE-seq data for patient BV-derived cells. Related to Figures 1, 2 and 3.**

(**A**) MultiUMAP generated with mRNA and AbSeq data from of 18941 leukemic cells isolated from murine BM colored by cluster (n = 3 mice). (**B**) Proportion of cells in the Pro-B, Large Pre-B and Small Pre-B stages of B-cell development per cluster. Pairwise comparison chi-square test with Bonferroni correction. (**C**) MultiUMAP colored by clone for each of the ten largest clones within the transcriptional landscape. (**D**) Normalised AbSeq expression across clusters. (**E**) Clone sizes per cluster. Each line represents a clone, colored by clone size. (**F**) Proportion of each clones (rows) in the two clusters (columns), clones are clustered with hierachical clustering. (**G**) Top 20 class predictive genes from XGBoost model ranked based on their prediction gain with their relative average expression (z-score) in each class in cluster 1. (**H**) Balanced accuracy with the Xgboost model (left bar) versus mean balanced accuracy from 1000 trainings with label permutations in the training set (right bar), error bar denotes standard deviation, p-value<0.05.

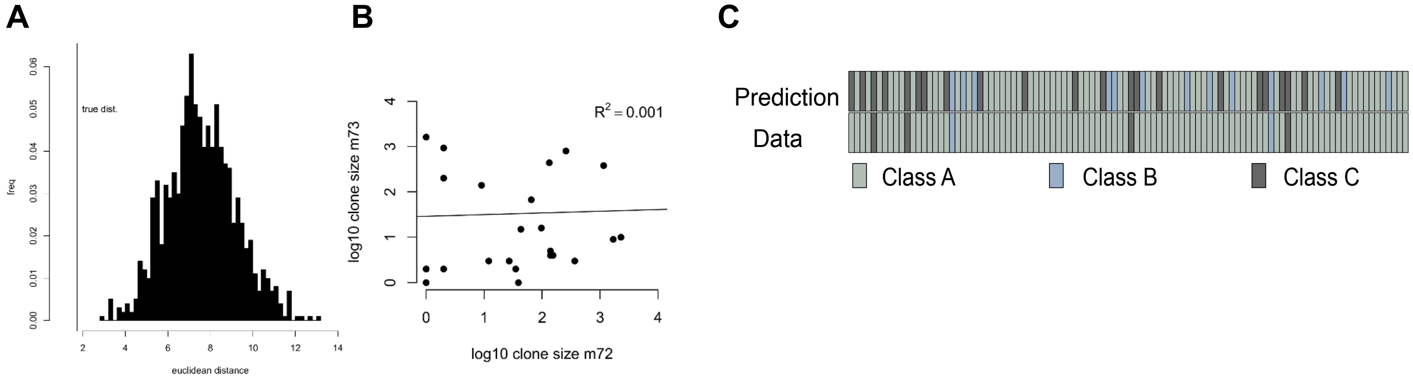


**Supplemental Figure 4. Comparison of patient CR-derived clone characteristics between mice for shared clones and differentiation class predictions. Related to Figure 3.**

(**A**) Euclidean distance between cluster distributions for the shared clones having at least 10 cells per mouse, in the dataset (vertical line) or for 1000 clone permutations (histogram). (**B**) Correlation of clone sizes for shared clones between mouse 1 (m72) and mouse 2 (m73). Line denotes linear regression. Values are log10 transformed. (**C**) XGBoost model prediction of differentiation class compared to experimental data for 100 randomly selected cells.

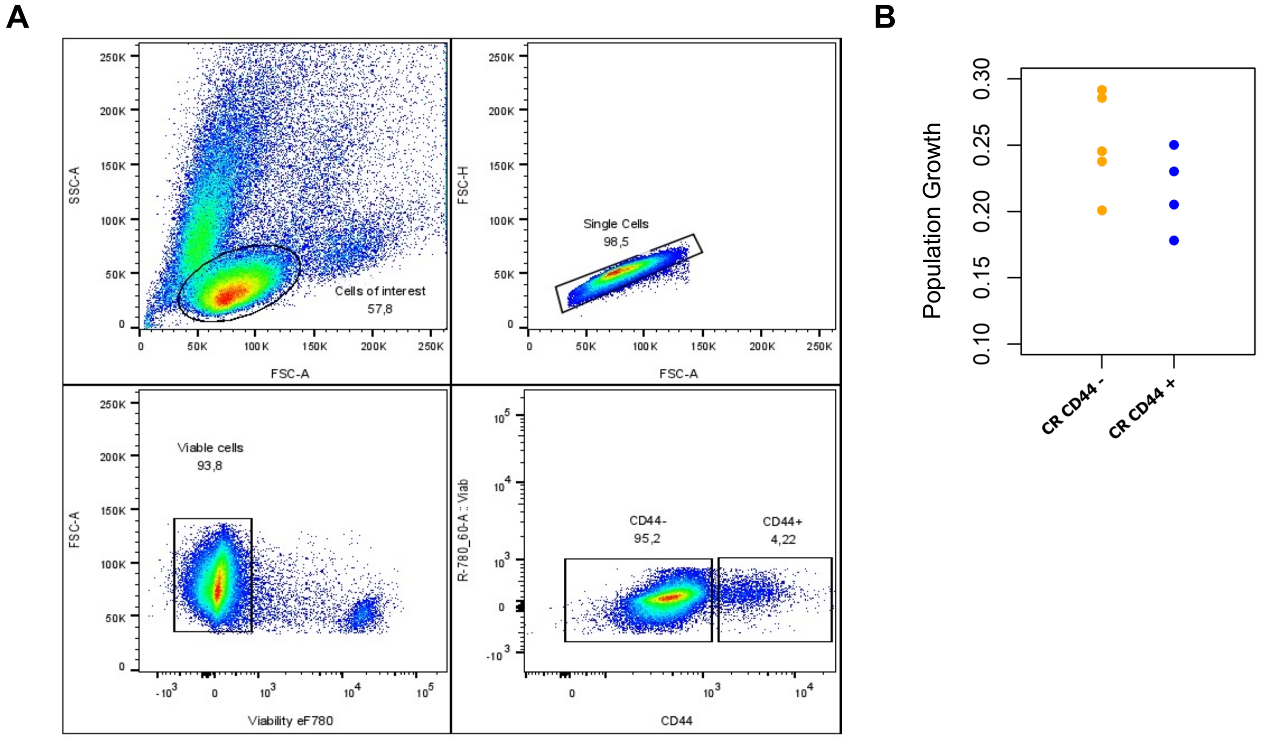


**Supplemental Figure 5. CD44+/- patient CR-derived B-ALL cells. Related to Figures 4**
(**A**) FACS gating strategy for prospective isolation of CD44+ and CD44- leukemic cells. (**B**) Population growth of CD44+/- CR cells *in vivo* (n=4-5 mice).

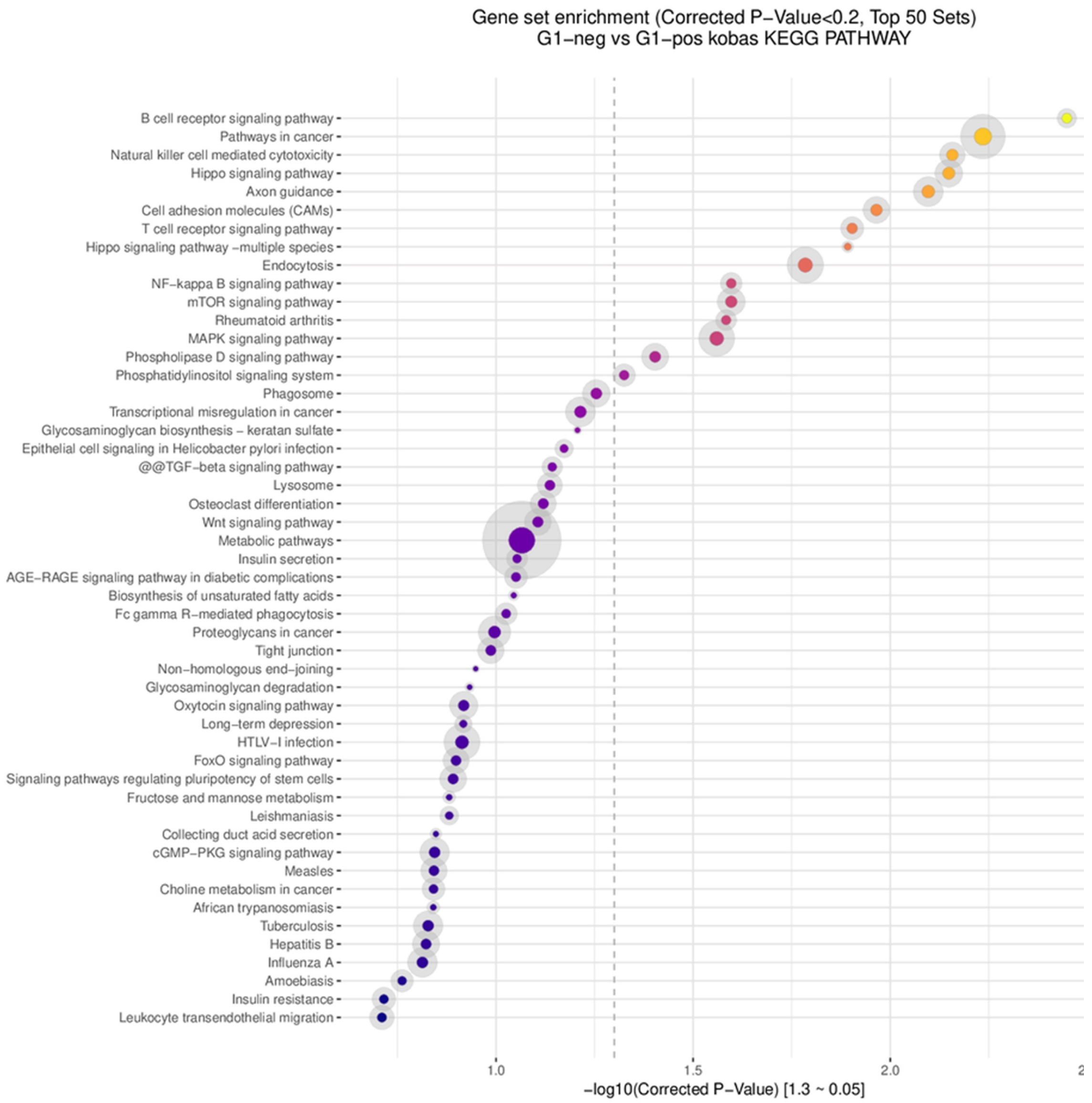


**Supplemental Figure 6. Pathway analysis.** Differential gene expression analysis of bulk RNA-sequencing of CD44+ and CD44- patient CR-derived cells after *in vivo* leukemogenesis reveals stronger B cell receptor signalling pathway (KEGG) activation in CD44+ cells.

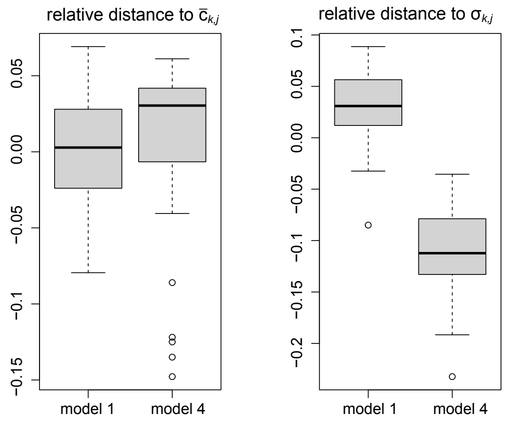


**Supplemental Figure 7. Patient CR, comparison of clone repartition across clusters between the dataset and 1000 simulations from posterior distributions**. Left panel: sums of the relative distances to the mean cell proportions per cluster per group of 6 clones $\overline{C}_{k,j}$ for model 1 (one-way differentiation with MHC-I cluster as source cluster) and model 4 (corresponding plasticity model). Right panel: sums of the relative distances to the standard deviation $\sigma_{k,j.}$ $\sigma_{k,j}$ represents the variation in cell proportions between clones for a given cluster and a given group (bin) of 6 clones while $\overline{C}_{k,j}$ is the corresponding mean.
